# Production, analysis, and safety assessment of a soil and plant-based natural material with microbiome- and immune-modulatory effects

**DOI:** 10.1101/2024.04.23.590687

**Authors:** Anirudra Parajuli, Iida Mäkela, Marja I. Roslund, Emma Ringqvist, Juulia Manninen, Yan Sun, Noora Nurminen, Sami Oikarinen, Olli H. Laitinen, Heikki Hyöty, Malin Flodström-Tullberg, Aki Sinkkonen

**Affiliations:** Center for Infectious Medicine, Department of Medicine, Karolinska Institutet, Karolinska University Hospital Huddinge, Stockholm, Sweden; Ecosystem and environment research program, Department of Ecological and Environmental, Science University of Helsinki, Helsinki, Finland; Horticulture Technologies, Natural Resources Institute Finland, Helsinki and Turku, Finland; Faculty of Medicine and Health Technology, Tampere University, Tampere, Finland

**Author notes:** Corresponding authors Aki Sinkkonen Horticulture Technologies, Natural Resources Institute Finland Turku, Finland, Malin Flodström-Tullberg, Center for Infectious Medicine, Department of Medicine Karolinska Institutet, Stockholm, Sweden. These authors contributed equally to the work (Malin Flodström-Tullberg, Aki Sinkkonen).

**Keywords:** Biodiversity hypothesis, environmental microbiota, microbial exposure, natural material, microbial diversity, soil

## Abstract

Reduced contact with the microbiota from the natural environment has been suggested to contribute to the rising incidence of immune-mediated inflammatory disorders (IMIDs) in the western, highly urbanized societies. In line with this, we have previously shown that exposure to environmental microbiota in the form of a blend comprising of soil and plant-based material (biodiversity blend; BDB) enhances the diversity of human commensal microflora and promotes immunoregulation that may be associated with a reduced risk for IMIDs. To provide a framework for future preclinical studies and clinical trials, this study describes how the preparation of BDB was standardized, its microbial content and safety assessments. Multiple batches of BDB were manufactured and microbial composition analyzed using 16S rRNA gene sequencing. We observed a consistently high alpha diversity and relative abundance of bacteria normally found in soil and vegetation. We also found that inactivation of BDB by autoclaving effectively inactivates human and murine bacteria, viruses and parasites. Finally, we demonstrate that experimental mice prone to develop IMIDs (non-obese diabetic, NOD, mouse model) can be exposed to BDB without causing adverse effects on animal health and welfare. Our study lays the foundation for a safe, sustainable, and affordable way to mimic exposure to natural microbiota that has the potential to have enormous health- and socio-economic impacts.

## 1. Introduction

The prevalence of autoimmune and inflammatory diseases, such as allergies, asthma and type 1 diabetes, has increased dramatically in recent decades, especially in western, highly urbanized countries [1,2]. Numerous studies have consistently revealed a direct relationship between atopic diseases and urbanization and western lifestyle, and the protective effect of farm environment and non-affluent traditional lifestyle against these diseases [3,4,5,6]. The biodiversity hypothesis proposes that exposure to natural environment such as vegetation and soil modulates the human microbiome, contributes to immune balance, and protects against allergy and inflammatory disorders [7]. This hypothesis is based in part on observations that the increase in the prevalence of immune mediated inflammatory diseases (IMIDs) correlates with a rapid loss of global biodiversity (reduction in biological species), which is probably mediated by reduced contact with natural microbial biodiversity [8]. Indeed, it has been suggested that disconnection of human with soil (and the microbiota they harbor) in an urban environment where soil is replaced by concrete and asphalts may be one of the reasons behind a compromised immune function and consequently a high incidence of immune-mediated disorders in the western urban society [9]. This will likely be an even more serious issue in the future since two-thirds of the world population is believed to live in urban areas by 2050 [10].

Microbes and microbial components are proposed to modulate the immune system through continuous activation of pattern recognition receptors (PRRs) on both skin and mucosal surfaces [11]. This phenomenon is particularly abundant in rural natural environments and less so in artificial urban settings. In support of this, studies have shown that exposure to a highly diverse environmental microbiota and microbial antigens rather than to a particular or selected microbial strain is beneficial in promoting immunoregulation and confer protection against immune-mediated disorders [12,13,14]. In addition, exogenous microbiota may also alter the composition of the commensal microbiota including that of the skin and gut in humans and animal models [15,16]. The human commensal microbiota, particularly the gut microbiota and their cellular components and metabolites, have a well-documented role in the development and function of the immune system [17,18]. In line with this, several studies have suggested that high exposure to soil microbes (and microbial components) and vegetation may be beneficial, and even necessary for the normal maturation of the immune system and protection against allergic and atopic diseases [19,20,21,22,23].

A few studies have explored exposure to soil and plant materials as a strategy to modulate the composition of commensal microbiota and immune function in animal models [16,24]. These studies have indicated that exposures can alter the composition of gut microbiota substantially and promote immune regulation. However, the exposures were carried out using one or very few soil or plant components. The microbial composition of soil depends on several environmental and anthropogenic factors [25]. This may lead to problems with reproducibility if the collection time and area as well as storage conditions of the soil and plants differ [26]. In addition, a particular soil product, such as sand or a mixture of sand and peat, may comprise of a limited composition of microbes [27]. Consequently, it may neither stimulate an adequate number of PRRs, nor contain sufficient number and richness of bacteria to produce immunomodulatory metabolites, resulting in limited immunomodulation. Likewise, the currently published studies [16,24] did not address the safety aspects of the exposure, which is of importance. Therefore, a broad range of microbiota from a variety of natural sources that can reproducibly stimulate a broad range of PRRs and provide immune modulatory metabolites in a safe manner is desirable [28].

We have recently manufactured a biodiversity blend (here denoted as BDB) from a variety of soil and plant-based materials as potential immunomodulatory extract. The goal was to produce a blend with high microbial biodiversity and density of non-culturable bacteria. In our previous studies we found that exposure to BDB altered skin and gut microbiota and immune status favourably in both children and adults [29,30,31]. These findings suggested that BDB may mimic nature exposure and that BDB will be a valuable tool to address the lack of such exposure more directly in the aetiology of inflammatory diseases. BDB may even be a promising candidate for the prevention of IMIDs. However, live BDB may pose a threat to immunocompromised people or experimental animal disease models since soil contains potential human and animal pathogens. Therefore, whether inactivation of live bacteria maintains the effect of BDB on microbial composition and immune system in exposed individuals is of interest when the BDB is to be considered in preclinical and clinical trials, particularly if intended to be used via routes of administration other than topical application e.g. as ingestables.

The composition of the BDB, its manufacturing process and analyses of microbial composition has not been described in detail. Likewise, the safety aspects of the BDB have also been not investigated thoroughly. Through this report, we describe the composition and manufacturing of multiple batches of BDB and how the production has been standardized. We also describe a way to effectively inactivate potential human and murine pathogens in the BDB so that it can be safely used in experimental animals and potentially immunocompromised humans. Finally, we addressed the safety of exposure to inactivated BDB in an immunocompromised mouse model.

## 2. Material and methods

### 2.1 Manufacturing of BDB

The BDB comprises of different forms of soil in varying proportions (referred to as major and minor components) and sphagnum moss (described in detail in the result section). All raw materials were composted and controlled according to EU regulations [30]. The raw materials were stored in containers covered by lids that were perforated with two holes containing thin pieces of cotton to allow aeration but to prevent fast drying [26]. The containers were stored in a cold room (5−7 °C) until mixed thoroughly together. The major ingredients were pre-sieved through a Ø 5 mm sieve size and the minor ingredient using a Ø 2 mm sieve. Moss was dried overnight in a tray dryer and then crushed and mixed thoroughly before mixing with other ingredients. Ultra-pure sterile milli Q water was poured into the mixture little by little until the soil got saturated. The soil water mixture was incubated at room temperature for 3 hours to allow the extraction of microbes into water phase as much as possible. The resulting extract was filtered through 125 µm sieves to collect water extract and smaller particles. The size of the sieve was chosen to ensure homogeneity of the filtrate without risk of clogging during the process. The flow through was collected in a new sterile bucket and transferred into sterile containers and frozen at -20°C overnight. The frozen material was then freeze dried at 0.25 bar pressure until complete removal of moisture. The freeze-dried material was stored in sealed containers at 5-7 °C.

### 2.2 Microbial composition analysis of BDB

Total DNA was extracted from 0.25 ± 0.07 g of BDB in triplicates, which were then subsequently pooled, using PowerSoil® DNA Isolation Kit (MolBio Laboratories, Inc., Carlsbad, CA, USA) according to the manufacturer’s standard protocol. Sterile water (250 µl) was used as a negative control. The quality of the extracted DNA was checked using agarose gel (1.5%) electrophoresis and quantified with Quant-iT™ PicoGreen® dsDNA reagent kit (Thermo scientific, MA, USA).

Sequencing of bacterial DNA from the BDB was done as described earlier [30]. Briefly, the V4 region within the 16S rRNA gene of bacterial DNA was amplified by PCR and sequenced using Illumina MiSeq platform (Illumina, San Diego, CA, USA). Raw sequences were processed following the Divisive amplicon denoising algorithm 2 (DADA2) pipeline (version 1.16) in the R environment (version 4.2.1) using the ’dada2’ package [32,33]. Quality profiles were generated for raw sequences, followed by trimming and filtering to remove low-quality bases, primers, and sequences with ambiguous bases using the ’filterAndTrim’ function.

Forward and reverse reads were merged using function ’mergePairs’. Dereplication was performed to reduce computational load by identifying and collapsing identical sequences. A sequence table was constructed and chimeric sequences were identified and removed using the ’removeChimeraDenovo’ function. Dereplication and denoising were utilized using DADA2 default parameters. Taxonomic classification was achieved using the SILVA reference database (version 138) [34] with the ’assignTaxonomy’ function. Sequences belonging to archaea, mitochondria and chloroplast, which accounted for 2.3% of the total sequences (53230 out of 2303697 sequences), were removed. The alpha diversity (Shannon index and observed species) as well as the relative abundance of bacterial taxa were visualized using ggplot2 function in R (v 4.3.2). The difference in community composition among all BDB batches were calculated using using permutational multivariate analysis of variance (PERMANOVA) and visualized through non-metric multidimensional scaling (NMDS) ordination using metaMDS and as dendrograms using ‘hclust’ functions in the vegan package (v2.6.4) in R [35]. Differences in the bacterial alpha diversity and relative abundance of bacterial taxa among the 11 batches of BDD was compared using analysis of variance (ANOVA) on normal or log and square-root transformed data and group-wise comparison was done using Tukey’s post hoc test. All p-values were adjusted using Benjamini-Hochberg method. Adjusted p-values <0.05 were considered statistically significant.

Raw sequences generated in the study are available in Sequence Read Archive in NCBI under accession number PRJNA1037346.

### 2.3 Inactivation of BDB

BDB (around 50ml) was transferred into individual autoclave bags under sterile conditions. Sealed bags were sterilized by autoclaving at 121°C for 40 min or at 134°C for 30 min.

Autoclaved BDB was stored at 4°C prior to microbiological analyses and usage.

### 2.4 Human pathogen analyses

Total nucleic acid (DNA and RNA) was extracted from BDB by PowerSoil DNA Isolation Kit or PowerSoil Total RNA Isolation Kit (Qiagen, Hilden, Germany). qPCR was carried out for viral and protozoan pathogens including enterovirus, rhinovirus, rotavirus, norovirus, Giardia and Cryptosporidium as described in Krogvold et al. [36]. A cut-off Ct value of 42 was used to identify positive samples. The qPCR tests were validated by spiking samples with known amount of target RNA. Spiked samples were analyzed by qPCR and results were compared to positive controls.

### 2.5 Rodent pathogen analyses

Genomic DNA was extracted from both autoclaved and non-autoclaved BDB using DNeasy Powersoil DNA extraction kit (Qiagen, Hilden, Germany) according to manufacturer’s standard instructions and quantified using nanodrop ND-1000 spectrophotometer (Thermo Fisher scientific, MA, USA). Total RNA was extracted from 50-100 mg BDB using TRIzol^TM^ reagent (Thermo Fisher scientific, MA, USA) according to manufacturer’s instructions and diluted in 50µL sterile MilliQ water. The BDB was homogenized at 15 Hz speed with 1.25 mm beads for 8 min in a Tissuelyzer (Thermo Fisher scientific, MA, USA) in 1mL TriZol prior to extraction. Extracted RNA was stored at -80°C until downstream analyses.

BDB (prior to or post autoclaving at 121°C or 134°C) was analysed for the presence of rodent pathogens according to the list of Federation of European Laboratory Animal Science Associations (FELASA) recommended quarterly check [37]. All testing was performed at a SPF testing accredited facility (https://felasa.eu/) at the National Veterinary Institute of Sweden, Uppsala, Sweden. Bacterial outgrowth from aerobic and anaerobic cultures were analysed with MALDI-TOF at the species level while DNA or RNA extracted form live or autoclaved BDB was screened for virus, parasites and bacteria by qRT-PCR (Table 1). The efficacy of pathogen inactivation was tested by autoclaving in standard (50 ml) or larger volumes of BDB and the reproducibility tested by autoclaving the BDB in three different runs.

**Table 1.** Species and genera analysed by PCR, RT-PCR or culture for the presence of murine pathogens.

| FELASA listed pathogens | Detection method |  |  |
| --- | --- | --- | --- |
|  | PCR, RNA | PCR, DNA | Culture |
| **Viruses** |  |  |  |
| Murine Hepatitis virus | X |  |  |
| Murine Rotavirus | X |  |  |
| Murine Norovirus | X |  |  |
| Murine Parvoviruses |  | X |  |
| Thelie's murine encephalomyelitis virus | X |  |  |
| **Bacteria** |  |  |  |
| Citrobacter rodentium |  |  | X |
| Clostridium piliforme |  |  | X |
| Corynebacterium kutscheri |  |  | X |
| Helicobacter spp (H. hepaticus, H. bilis and H. typhlonius) |  | X |  |
| Mycoplasma pulmonis |  | X |  |
| Rodentibacter heylii |  |  | X |
| Rodentibacter pneumotropicum |  |  | X |
| Salmonella spp. |  | X |  |
| Staphylococcus aureus |  | X |  |
| Streptobacillus moniliformis |  |  | X |
| Streptococci beta haemolytic (not group D) |  |  | X |
| Streptococcus pneumoniae |  |  | X |
| **Endoparasites** |  |  |  |
| Spironucleus muris |  | X |  |
| Enterobius spp. |  | X |  |
| **Exoparasites** |  |  |  |
| Myobia spp. |  | X |  |
| Rhodfordia spp. |  | X |  |
| Myocopte spp. |  | X |  |

### 2.6 Inactivated BDB exposures in mice

A specific-pathogen-free (SPF) colony of inbred male and female non-obese diabetic (NOD) mice aged three weeks were randomly assigned to exposure or control groups. NOD mice are prone to develop autoimmune type 1 diabetes at an age above 10-12 weeks [38,39]. Diabetes incidence is high in female mice (around 60-90% at an age of 6 months) but low in males (around 10-25% at an age of 6 months). We initially performed a short-exposure duration (3 weeks) in young nondiabetic male mice and, when the treatment proved safe, explored the same but longer exposure (6 weeks) in young non-diabetic female NOD mice. Mice in the exposure group were exposed to BDB autoclaved at 134°C and the mice in the control group were mock exposed to the cage bedding material alone. Exposures were carried out in an SPF accredited facility and mice were allowed to acclimatize to the exposure room for a few days before the start of exposure. Exposure to BDB was carried out by sprinkling one pack of 50mL autoclaved BDB over cage bedding material in a new cage. Mock exposures were always carried out before BDB exposures, and exposures were carried out at the same time of day throughout the study. Mice from a single cage (maximum of 5 in each cage) were transferred to the cage containing the BDB or bedding alone and were allowed to play freely for 30 min after which they were transferred back to their original cage. A new (unused) cage was used for each exposure for both BDB and mock exposure. The cages were opened, and animals handled inside a laminar hood. The exposure dose was chosen based on our previous observation that this amount of BDB induced a visual exposure effect such as presence of the BDB in body parts including fur, tail, nose, mouth, and ear [40]. Male NOD mice were exposed to BDB (n=5) or mock (n=5) five times a week on weekdays for three weeks and were left unexposed for a further six weeks to monitor the (long term) effect of BDB exposure after exposure was discontinued. Female NOD mice were exposed to the BDB (n=9) or mock (n=8) five times a week on weekdays for six weeks. All animals were fed with the same diet and water ad libitum and were kept in the same room in an individually ventilated cage (IVC) system. Animal health was monitored by weekly weighing as well as through FELASA scoring that take into account different parameters including visual inspection of animal status and behavior (Supplementary information 1). Blood glucose level was measured for all mice from tail vein blood at the experimental endpoint to assess whether any of them had developed diabetes as described earlier [39].

Ethical approval for all animal work was obtained from Stockholm Southern Animal Ethics Board. Mice were housed and maintained in accordance with EU Directive 2010/63/EU for animal experiments.

#### 2.6.1 Pulmonary health assessment

At experiment endpoint, mice were euthanized using isofluorane followed by cervical dislocation. Whole lung was fixed in 4% PFA over-night, then transferred to 70% ethanol until being dehydrated and embedded in paraffin. Single 5 µM sections were stained with hematoxylin and eosin and scored by a researcher blinded to treatment groups. Overall health of the pulmonary structure was based on level of perivascular and peribronchial inflammation, fibrosis, parenchymal and alveolar damage, and red blood cell infiltrate.

Ordinal degree scores were set on total over all detection of unhealthy morphology (0–4; absent, rare, multifocal, coalescing, diffuse) as described previously [41].

## 3. Results

### 3.1 Material composition of BDB

To create a nature-derived material mimicking wild nature of the Nordic countries and containing a very high diversity of microbiota we included a variety of soil and plant materials harboring a diverse range of microbiota. The components of the BDB were chosen after careful consideration of microbial richness and diversity ease in availability and the presence and abundance of potentially pathogenic microorganisms. The components consist of leaf compost, industrially manufactured agricultural soil and compost made of forestry and agricultural side streams in equal proportion (29.6% each). These together constitute 89% of the BDB. *Sphagnum* moss, the dominant moss of Nordic countries (https://artfakta.se/naturvard/taxon/1004718), was added to a final constituency of 7.4%.

Minor components added were potting soil, meadow soil, lawn soil, infrastructure/park soil and gardening soil at 0.7% each (Table 2). The difference in the major components and minor components was primarily the microbial diversity with the major components having a higher diversity compared to the minor components [26]. As previously reported, there were only minor variations in microbial composition, dry weight, organic matter, pH, soil characterization, nutrients (NH4^+^, No3^-^, PO4^-^) and elements including Cd, Al, Co, Cr, Cu, Mn, Ni, Fe, Zn, P, As, Pb before and during the storage time [26].

**Table 2.** Material composition of biodiversity blend.

| Material | Trade (Finnish)<br>name | Percentage<br>(volume) | Supplier |
| --- | --- | --- | --- |
| **Major ingredients** |  |  |  |
| Leaf compost | Lehticomposti | 29.6 | Kekkilä, Finland |
| Agricultural soil | Maatalousmaa | 29.6 | Kekkilä, Finland |
| Forestry-agricultural side<br>stream compost | Metsä-<br>maatalouden<br>sivuvirtakomposti | 29.6 | Biolan, Finland |
| **Sphagnum moss** | Sammal | 7.4 | Biolan, Finland |
| **Minor ingredients** |  |  |  |
| Potting soil | Musta multa | 0.7 | Biolan, Finland |
| Meadow soil | Niittymulta | 0.7 | Kekkilä, Finland |
| Lawn soil | Nurmikkomulta | 0.7 | Kekkilä, Finland |
| Infrastructure/park soil | Inframulta | 0.7 | Kekkilä, Finland |
| Gardening soil | Viljelymulta | 0.7 | Kekkilä, Finland |

### 3.2 Formulation of a powder like product

We next set out to create a powder-like consistency of the BDB for easy application and formulation into topical products. Therefore, the soil components were first dried, sieved or crushed (in case of moss) to facilitate the mixing process (see materials and methods section for detailed information). Water extraction filtration was carried out to retain most of the microbiota without the presence of larger components of soil and moss. The filtrate was then freeze-dried to allow the microbes to remain in the BDB powder. Eleven batches have been produced until now using this method (with few variations as listed in Supplementary table 1) and analysed in detail.

### 3.3 Microbial composition of the BDB

The manufactured BDB batches were analysed for their microbial composition. DNA was extracted and the composition was analysed using 16S rRNA gene sequencing as described in the materials and methods section. The microbial composition of all individual raw material components has been reported before [26] and the microbial composition of one of the lots of batch 10 was already described in Nurminen et al. [29]. PERMANOVA revealed that the bacterial community composition was different among the 11 batches p=0.001). All the batches had a high diversity measured as Shannon’s index, ranging from 5.35 to 6.6 (median) and observed species ranging from 280 to 1200 (Figure 1). Major differences in the diversity and richness were observed between the first two and last batches compared to the rest of the batches (Supplementary Table 2). Batch 8 had slightly higher intra-batch variation in Shannon’s diversity index and richness (Figure 1) but was within the overall variation among all BDB batches. Hierarchical clustering revealed that different lots of a single BDB batch clustered together with similar microbial composition for the first five batches compared to the last 6-11 batches (Figure 2a). This was also observed in the NMDS, where the bacterial community composition of the first five batches were different from the subsequent seven batches produced (Figure 2b). Proteobacteria was the most abundant phylum in all batches ranging from 31% (batch 1) to 37.5% (batch 11) of the BDB followed by Bacteroidota (14- 22%) and Chloroflexi (8-14%) (Figure 3). A high variation in the relative abundance of bacterial phyla such as Proteobacteria, Bacteroidota and Acidobacteriota was observed among the BDB batches whereas less variation was observed for the relative abundance of Chloroflexi, Actinobacteriota and Verrucomicrobiota (Supplementary Table 2).

**Figure 1:**
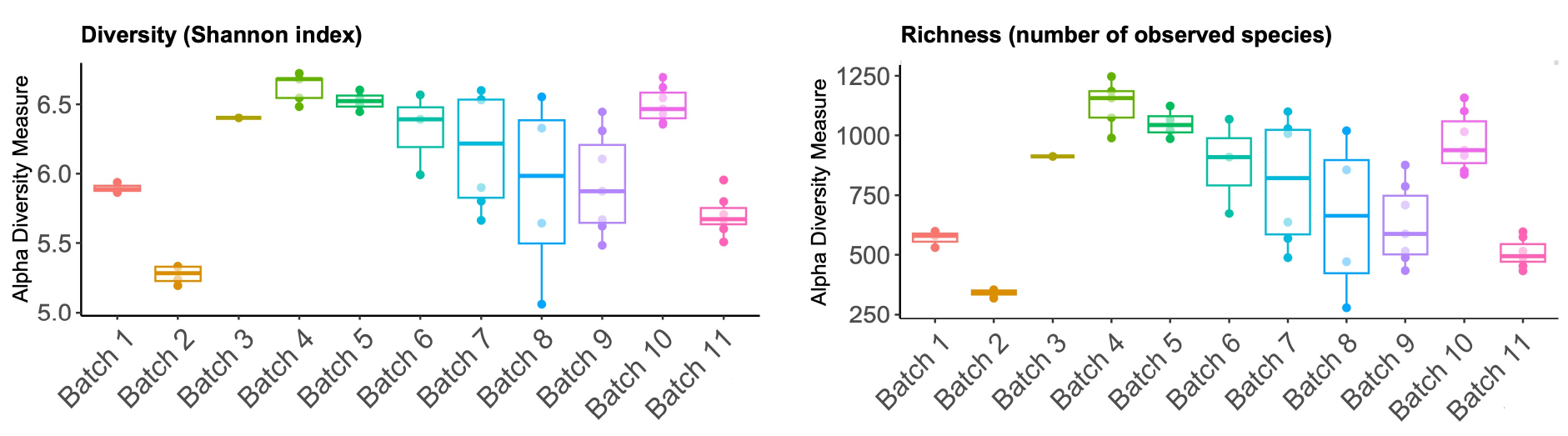
BDB contains a high diversity and richness of bacteria. 16S rRNA sequencing was performed on DNA extracted from 11 batches of freshly produced BDB and alpha diversity of the identified bacteria was assessed using amplicon sequence variants (ASVs) from the sequence data. The diversity (Shannon index) and richness (number of bacterial species) were found to be consistently high in all 11 batches. The points in each box plot represent different lots of BDB sampled within the batch. Statistics are shown in Supplementary Table 2.

**Figure 2.**
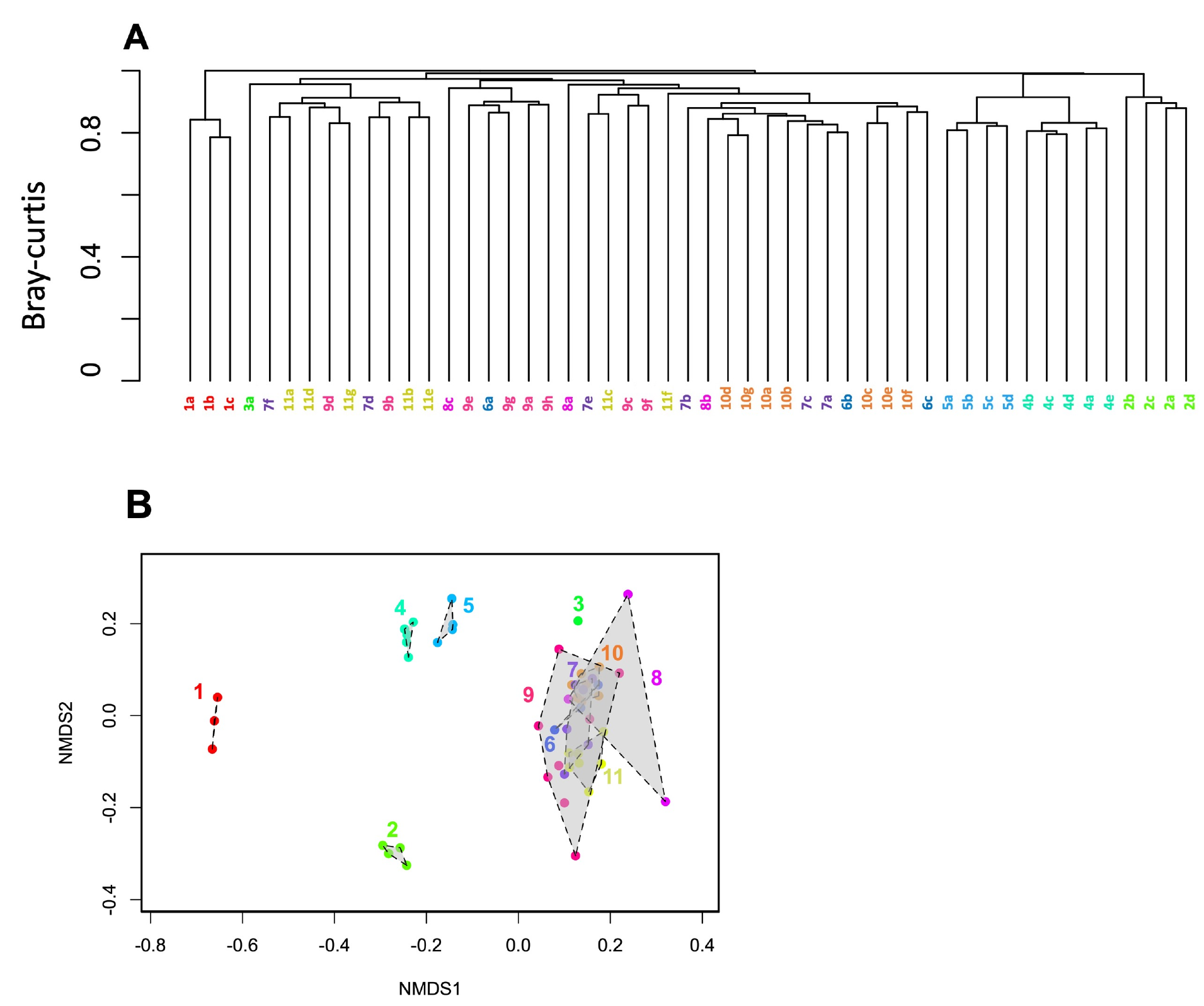
The community composition of bacteria in the BDB batches. A. Hierarchical clustering based on group average linkage using bray-curtis dissimilarity metric was done for all samples analysed from 11 BDB batches using ‘hclust’ function in vegan package in R. The dendrogram is based on the relative abundance of ASVs and reveals the similarity or dissimilarity in ASV distribution across 11 batches of BDB. The alphabets represent the lots sampled within the batch. Batch one outgroups from the others, and a second bifunction is seen between batch 2-5 and 6-11. **B**. Non-metric multi-dimensional scaling (NMDS) was done to visualize the community composition of bacteria within and between the batches based on relative abundance of ASVs using bray-curtis dissimilarity metric using metaMDS function in vegan package in R. The numbers represent the batches.

**Figure 3:**
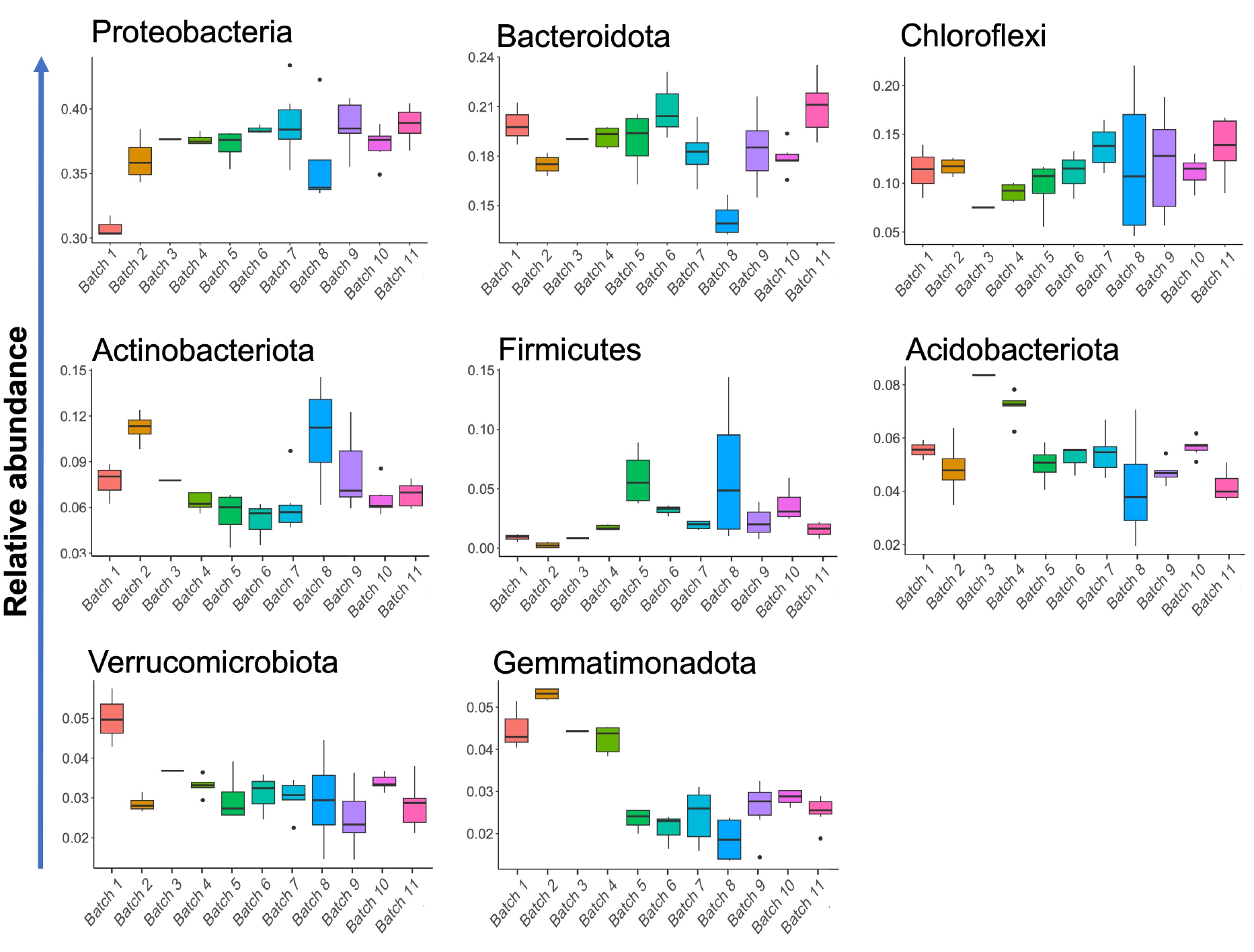
The BDB comprises of a high abundance of bacterial phyla normally found in environmental samples such as soil and plants. The relative abundance of bacterial phyla from the 16S rRNA sequencing data was compared among the 11 batches of BDB. Phyla representing more than 1% of the total bacteria are shown. The relative abundance of the most abundant bacteria was different whereas those of less bacteria were similar among the BDB batches (Statistics in Supplementary Table 2).

### 3.4 Inactivation of BDB

We next set out to neutralise any potential pathogens in the BDB to create a sterile product. To this end, we selected one BDB batch (batch 10) and autoclaved four individual samples from this batch at 121°C or 134°C, extracted DNA and RNA, and compared the presence of pathogens in autoclaved and untreated BDB.

We first analysed the presence of pathogenic human viruses and parasites commonly present in soil, namely adenovirus, enterovirus, rhinovirus, norovirus, sapovirus, parechovirus, influenza A virus, hepatitis A virus, Cryptosporidium and Giardia. We did not detect any of these pathogens even in the live BDB (Supplementary Table 3).

To enable the use of BDB in pre-clinical trials, the material was analysed according to the standardized FELASA specific pathogen list [37] (Table 1). We assessed the presence of rodent pathogens including bacteria, viruses, mites, and helminths through aerobic and anaerobic culture of autoclaved BDB as well as PCR of DNA and RNA extracted from both live and autoclaved BDB (details in material and methods). In BDB autoclaved at 121°C, aerobic outgrowth of environmental bacteria *Fictibacillus spp., Micrococcus luteus, Macrococcus brunensis, Neisseria subflava*, and two murine pathogens *Bacillus megaterium,* and *Staphylococcus spp.* was detected, while no bacterial outgrowth was detected in the anaerobic culture (Table 3). Likewise, Parvovirus DNA and pinworm (*Enterobius vermicularis*) DNA were detected through PCR analyses in both live and the BDB autoclaved at 121°C. We observed that after autoclaving at 134°C, there was no outgrowth of any environmental or pathogenic bacterial species nor the detection of bacterial, viral or parasitic nucleic acids (Table 3 and Supplementary Table 3). The reproducibility of the inactivation at 134°C was confirmed by autoclaving additional BDB samples also in varying amounts (data not shown).

**Table 3:** Comparison of presence of murine pathogenic bacteria, virus and parasites in live and BDB autoclaved at 121 °C or 134 °C.

|  | Bacteria |  |  | Virus | Parasites |
| --- | --- | --- | --- | --- | --- |
| Detection method | Aerobic culture | Anaerobic culture | Nucleic acids | Nucleic acids | Nucleic acids |
| **Live BDB** | Not tested | Not tested | None detected | Parvovirus DNA | *Enterobius vermicularis* DNA |
| **BDB autoclave d at 121°C** | *Bacillus megaterium* and *Staphylococcus spp.* | None detected | None detected | Parvovirus DNA | *Enterobius vermicularis* DNA |
| **BDB autoclave d at 134 °C** | None detected | None detected | None detected | None detected | None detected |

### 3.5 Sterile BDB is safe to use in an immunodeficient experimental mouse model

To assess whether exposure to inactivated BDB is safe in an experimental animal model prone to develop an immune mediated disorder, we exposed young (3 weeks of age) male and female NOD mice to BDB or in mock as described in materials and methods section. We monitored the effect of exposure to BDB on several health parameters including weight gain, animal appearance and behaviour. We observed that 30 minutes weekly (5 times per week) exposure to BDB was well-tolerated. Notably, the mice did not only move around in the BDB but also consumed part of it. We observed no adverse impact on weight gain during the exposure period or after the exposure was discontinued (9 weeks of age, Figure 4). None of the animals developed diabetes (data not shown). Pulmonary health of the animals was assessed at the endpoint and did not reveal any noticeable differences in pulmonary health between exposed or mock exposed animals. We also investigated signs of inhaled particulate matter but did not detect any (data not shown).

**Figure 4:**
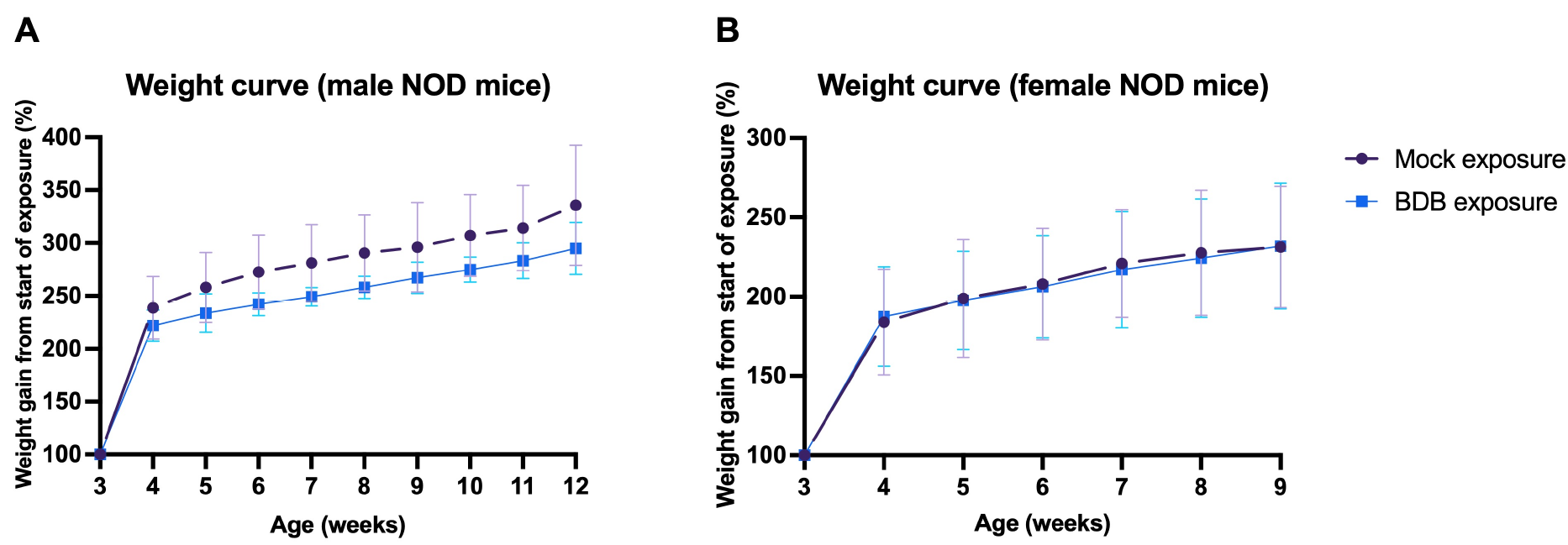
BDB exposure had no adverse effect on weight gain in NOD mice. (**A**) Male NOD mice (age 3-4 weeks) were exposed to mock (n=5) or BDB five times a week (n=5)) followed by six weeks of observational period. (**B**) Female NOD mice (age 3-4 weeks) were exposed to mock (n=8) or BDB five times a week (n=9) for 6 weeks. Weight measurements were done weekly and shown relative to the weight at 3 weeks. Data is presented as percentage weight gain, mean ± SEM.

## 4. Discussion

In this report, we describe the production of a blend with potential microbiome- and immune modulatory capacity. BDB was produced from commercially available soil and plant-based material, each with a well-characterized microbial composition [26] and in a well-defined proportion. Commercially available soil and plant-based materials that are accessible with relative ease from natural resources in Finland were used as the components of BDB. Soil is a large natural reservoir of microbial diversity whose beneficial effects on human health through direct and indirect means have been well documented [42,43,44]. Soils and plants from Northern Europe, particularly protected areas and other natural and seminatural ecosystems, are relatively free from typical anthropogenic disturbances, such as pollution, irrigation, fertilization, invasive species, human trampling and droppings of domestic animals [45]. They are thus close to being microbiologically and chemically pristine [46], which is why organic materials from those areas were chosen as the ingredients of the BDB. However, natural phenomena such as seasonal variation, moisture, sunlight, soil pH, nutrient contents as well as anthropogenic disturbances such as deforestation, urbanization, land use change, agriculture etc. alter the soil composition and microbiota considerably [25,43,47]. Therefore, we acknowledge that the BDB produced in places that differ in terms of climate and land use may not be microbiologically identical even using our established protocol.

We undertook a series of steps to produce a powder-like consistency that provides a flexibility of incorporating the BDB into a broad range of formulations intended for human clinical or animal (pre-clinical) trials. We have previously evaluated the efficacy of the BDB in modulating the composition of commensal microbiota and immune functions via topical application through hand rubbing that was mostly suitable for adults [29,30] or by mixing in sandboxes aimed for daycare children [28]. We observed that topical exposure was safe and could modify not only the skin microbiota but also potentially gut microbiota probably through (accidental) ingestion as well as inhalation, which is more probable with the powder like consistency of BDB.

We observed that the BDB had a very high number of bacterial species (richness) and diversity. However, there was some inter-batch variations in the alpha diversity as well as the relative abundance of bacterial phyla. This could be due to insufficient blending or the storage time of the ingredients since the production technique and storage conditions remained unaltered. We have previously shown that long term storage of the components of BDB alters the relative abundance of several bacterial taxa [26]. Indeed, we observed that there was a high variation in the relative abundance of major bacterial phyla (Figure 3) among the BDB batches compared to others although the diversity and richness were not compromised to a great extent. Our previous study showed that long-term storage led to changes in bacterial community composition in the raw materials of BDB [26]. This could also partly explain the variation in the relative abundance of bacterial phyla in the BDB. This could perhaps raise concerns about the reproducibility in modulating the microbiome or immune function among different batches of BDB owing to this difference in the relative abundance of bacterial taxa. However, while some studies have reported associations between exposure to certain bacterial taxa or groups and immune function and overall health [48,49], there is increasing evidence to suggest that exposure to diverse microbes and microbial antigens rather than one or few microbial strains could modulate the immune function more efficiently [12,13,14]. It is therefore reasonable to assume that small alterations in the relative abundance of bacterial taxa present in the BDB may not compromise the efficacy of the BDB as long as the diversity and richness remain high. We believe that the BDB having such high diversity and richness could be a simple yet very effective way to enhance microbial exposure, especially among urban inhabitants who are deprived of adequate microbial exposure due to urbanization and insufficient contacts with natural biodiversity[21,50]. Noteworthily, we have always used dry BDB. If autoclaved soil is wetted, it may serve as an effective inoculum for microbes, and hence any potential use of moist BDB needs several safety tests before applied in a clinical trial.

We acknowledge that live BDB may contain potential pathogens. This is not surprising since soil in general harbors a vast variety of microbes, many of which may be potentially pathogenic. However, in our own previous studies, we observed that exposure to BDB did not induce adverse health effects in healthy individuals but instead seemed to promote immunoregulation [28,29,30,31,51]. In fact, children in daycare centers are in daily contact with many sources of potential pathogens, including those carried by other children and other present in daycare yards. The added exposure to BDB in, for example sandboxes was not associated with infections or other side effects [20,22,28,52]. This is in line with reports suggesting that exposure to soil pathogens can in fact promote immune tolerance and a better functioning immune system [43,45,53]. In this context, is may be of relevance that the manufacturing process of BDB involved a careful mixing that effectively homogenizes the blend and thus breaks clusters of conspecific bacteria, including potential pathogen.

Furthermore, we always used the BDB in freeze-dried form that prevents outgrowth of any potential pathogens that could be present in low relative abundance. In any event, in any future study using the live BDB, we recommend assessment of safety in the same way as it has been measured in the previous studies.

The use of live BDB could, however be a problem if it is intended to be used in immunocompromised and/or a chronically ill population. Exposure to even less virulent pathogens may cause opportunistic infections in such individuals [54,55], which also applies to animals, particularly disease models of immune-mediated disorders. In addition, the topical application of live BDB may pose a problem in subjects with broken skin barrier such as cuts and wounds. A potential solution to this is to use sterile BDB. We therefore inactivated the freeze-dried BDB and observed that autoclaving at 134 °C was effective in removing known human and murine pathogens and did not induce any adverse health effects in the experimental mice that were exposed to or even consumed the BDB. The dose (50 ml) of BDB, frequency of exposure (5 times a week) and duration (30 min per exposure for six weeks) represent arbitrary exposure regimen and it remains to be seen whether autoclaving followed by such treatment alters the potential of BDB to alter skin and gut microbiota, stimulate PRRs and promote the production of metabolites that can modulate immune function. A chemical analysis of sterile BDB could provide information on its content of immunoregulatory components. Notably, bacterial endotoxins, including lipopolysaccharides (LPS), that are major antigenic determinants of gram-negative bacteria are resistant to autoclaving and heat treatments [56]. In addition, our previous study revealed that BDB autoclaved at 121°C exhibited immunoregulatory potential in C57BL/6 mice and stabilized the microbiome [40]. However, we here observed that autoclaving at 121°C does not effectively remove all murine pathogens suggesting that autoclaving at 134°C is a better way to sterilize the product.

We observed that proteobacteria was the major bacterial phyla in all BDB batches we have produced so far. Taxa belonging to Proteobacteria are known producers of endotoxins, which could potentially raise concerns about the potential pathogenicity associated with the use of BDB. However, Protoebacteria is also a major component of skin microbiota and is associated within innate and adaptive immune response. Several studies have shown that a higher relative abundance of skin Protoebacteria and several taxa within it is associated with immunomodulation at local (skin) or systemic level [13,20,57,58]. Importantly, we have observed in several of our studies that exposure to non-inactivated (live) BDB is safe in children and healthy adults without any reported pathogenicity [20,29]. This was observed in animal studies where exposure to inactivated or even non-inactivated BDB was well tolerated without any adverse health issues [40,59]. Moreover, lipopolysaccharides (LPS), which are the endotoxins that are produced by Proteobacteria are known immunomodulatory molecules [60]. Animal studies have also demonstrated that LPS treatment of mothers shape immune system development and anti-inflammatory activities in offspring [61,62]. As such, there is ample evidence to suggest that higher abundance of Proteobacteria in the BDB is not necessarily bad and may actually be beneficial.

A limitation of our study is that we analyzed the bacterial composition of the BDB in detail and no other microbes. Fungi, viruses and other microbes could also play a role in modulating human commensal microbiota composition and induce immune regulation, and as such their composition in BDB may be explored in future studies. Bacteria are however the most well-studied microbes in terms of immunomodulation and therefore we focused on bacteria in our current study.

## 5. Conclusions

In this report, we describe the production, microbial analyses and safety aspects of a soil and plant derived natural product, which has been shown to have microbiome and immune modulatory capacity in previous studies. We showed that the alpha diversity of environmental bacteria remained consistently high among all the BDB batches produced with some alterations in the relative abundance of some bacterial taxa. We also showed that the BDB was free of specific human pathogens and inactivation by autoclaving at 134°C inactivated any potential human as well as murine pathogens. The powder-like form of the BDB allows a flexibility to be incorporated into a wide range of formulations for different routes of administration for potential preclinical and clinical trials. Apart from the topical applications described before, BDB has further been combined with lotions intended for topical use in infants (Clinical trial: NCT03872219) and can similarly be used in cosmetic products. Depending on the outcomes and intended use and dose, the BDB can further be formulated into consumable formulations such as capsules. Likewise, in animal studies, we found that the BDB was easy to use by sprinkling a freeze-dried material over the cage bedding material. The exposure was well tolerated by immunocompromised mice in topical exposure through the cage bedding, potential inhalation and even consumed without development of side-effects. Future studies as needed to optimize the frequency, duration and dose of exposure for optimal benefits in terms of gut microbiota composition or immunoregulation. Use of highly sustainable and safe natural ingredients for the prevention and/or treatment of immune-mediated health disorders will likely have great medical and economic impact in the future.

## Acknowledgements

We thank the Institute for Molecular Medicine Finland (FIMM) and Environmental Laboratory AlmaLab for the laboratory analyses, and the IT Center for Science Finland (CSC) for computational services. We would also like to thank Ms. Selina Parvin, Karolinska Institutet, Stockholm, Sweden, for her help with the histological analyses of mouse lung sections, Dr. Virginia Stone for scientific discussions and assistance with mouse-exposure studies, the animal staff at the Preclinical Laboratory (PKL) Facility at Karolinska Institutet, Stockholm, Sweden, for their assistance with the mice studies.

---

**Supplementary Table 1.** Inter-batch variation in the manufacturing process of BDB.

| **BDB batch** | **Major ingredient same as table 1 (Yes/No)** | **Minor ingredient same as table 1 (Yes/No)** | **Date of manufacture** | **Deviation in manufacturing process** |
| --- | --- | --- | --- | --- |
| 1 | Yes | Yes | 28.08.2018 |  |
| 2 | Yes | Yes | 07.02.2019 |  |
| 3 | Yes | Yes | 14.02.2019 |  |
| 4 | Yes | Yes | 21.02.2019 |  |
| 5 | Yes | No | 07.03.2019 | Different kind of lawn soil used |
| 6 | Yes | Yes | 13.03.2019 |  |
| 7 | Yes | Yes | 21.03.2019 |  |
| 8 | Yes | Yes | 27.03.2019 |  |
| 9 | Yes | Yes | 01.04.2019 |  |
| 10 | Yes | Yes | 06.04.2019 |  |
| 11 | Yes | Yes | 15.04.2019 |  |

**Supplementary Table 2.**
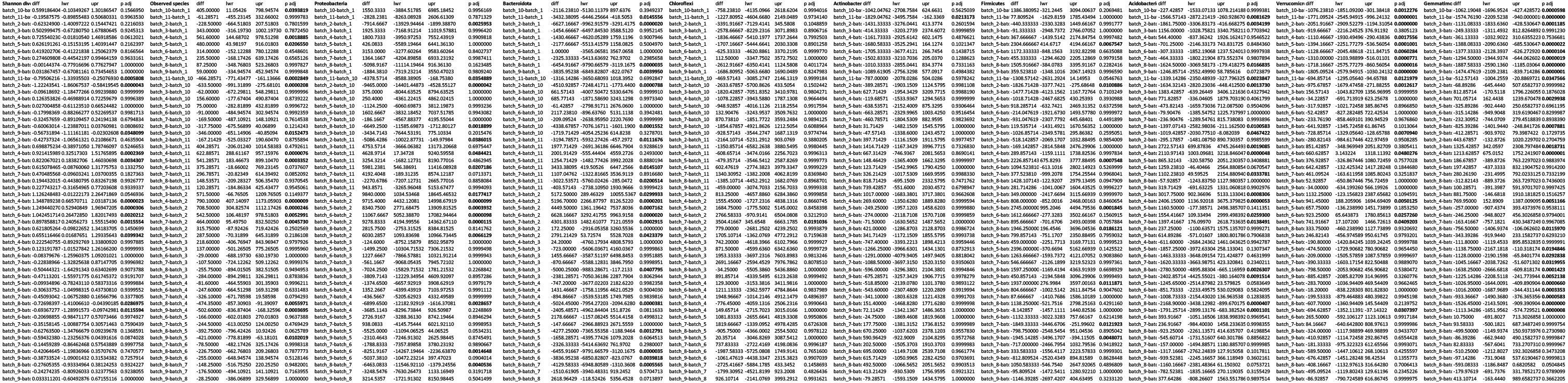
Tukey’s pairwise comparison for the the alpha diversity of overall bacteria and relative abundance of major bacterial phyla among the 11 batches of BDB Values in bold represent adjusted p values <0.05.

**Supplementary Table 3.**
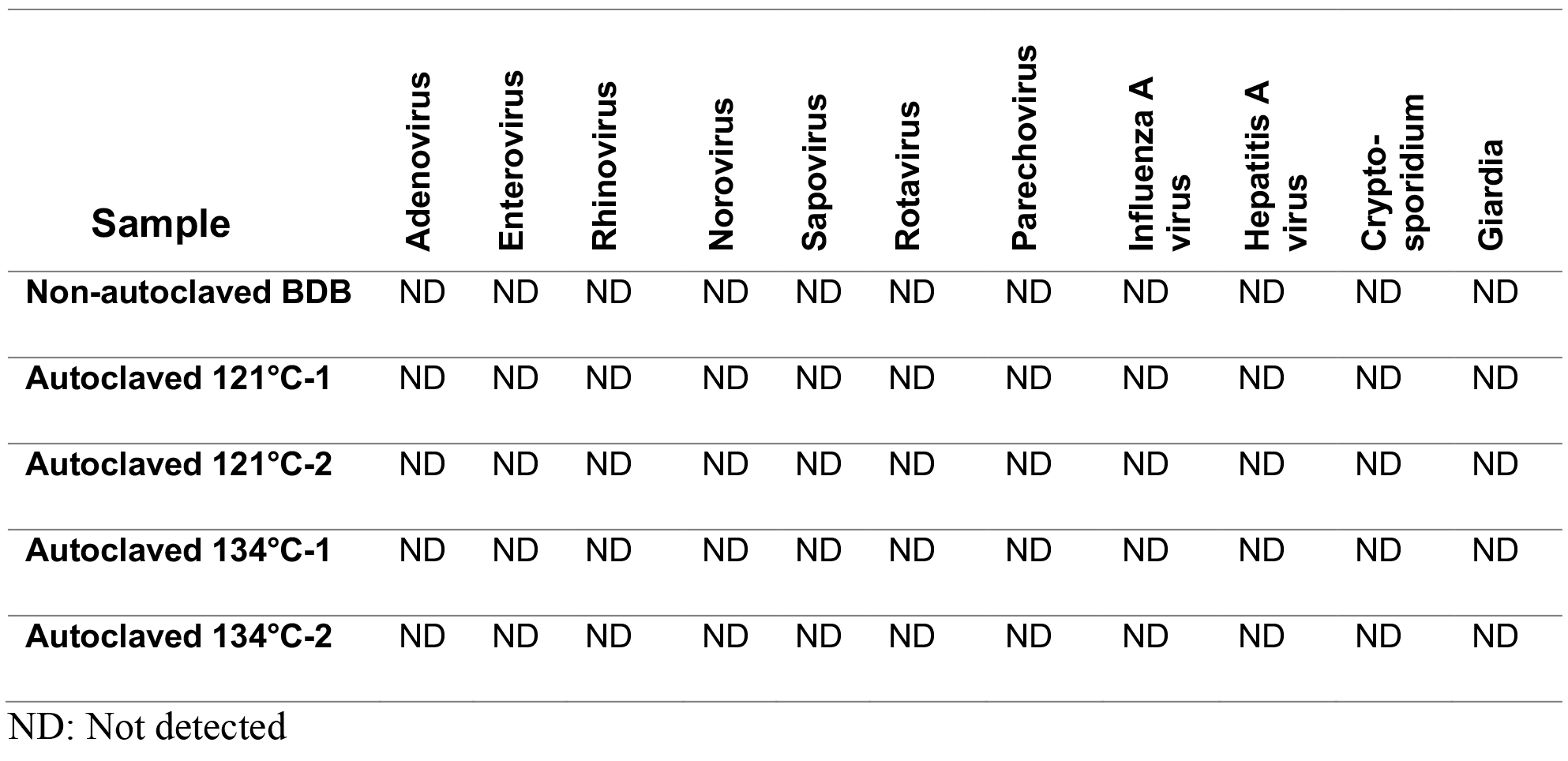
Common soil pathogens tested for their presence in the live (Non-autoclaved) BDB and BDB inactivated by autoclaving at 121°C (n=2) or 134°C (n=2).

| Sample | Adenovirus | Enterovirus | Rhinovirus | Norovirus | Sapovirus | Rotavirus | Parechovirus | Influenza A virus | Hepatitis A virus | Crypto-sporidium | Giardia |
| --- | --- | --- | --- | --- | --- | --- | --- | --- | --- | --- | --- |
| Non-autoclaved BDB | ND | ND | ND | ND | ND | ND | ND | ND | ND | ND | ND |
| Autoclaved 121°C-1 | ND | ND | ND | ND | ND | ND | ND | ND | ND | ND | ND |
| Autoclaved 121°C-2 | ND | ND | ND | ND | ND | ND | ND | ND | ND | ND | ND |
| Autoclaved 134°C-1 | ND | ND | ND | ND | ND | ND | ND | ND | ND | ND | ND |
| Autoclaved 134°C-2 | ND | ND | ND | ND | ND | ND | ND | ND | ND | ND | ND |

ND: Not detected

**Supplementary information 1**. General health monitoring of experimental mice The general health of the mice was observed and noted down throughout the study period so that any deviations from the normal microbiota composition could be tracked. The general health monitoring included:

Weight measurements, health status:

- General condition:

`o` Awake, reacts to stimuli, active
`o` Burrows/hides/lies still/startled when handled
`o` Immobile/little no movement/very afraid or aggressive
- Movement and posture:

`o` Normal
`o` Hunched, moderate lack of coordination
`o` Very uncoordinated/hunched posture and or back
- Piloerection:

`o` Fur smooth and well-groomed
`o` Moderate piloerection
`o` Severe piloerection, sticky and poorly groomed fur
- Skin:

`o` Completely covered with fur
`o` Small sores/scabs, scratching (or signs of), no infection
`o` Signs of infection (redness, pus), sticky and poorly groomed fur
- Weight:

`o` <5% weight loss compared to weight at day 0
`o` 5-10% weight loss compared to weight at day 0
`o` >15% weight loss#, compared to weight at day 0
- Porphyrin staining:

`o` No discolouration, clean and clear eyes
`o` Some porphyrin and/or discharge around the eyes and nose
`o` Obvious porphyrin on face/legs/paws
- Respiration:

`o` Breathing normal
`o` Breathing with open mouth, wheezy, panting
- Appetite:

`o` Normal, eating dry food, food disappears
`o` No signs animal has eaten dry food

No signs animal has eaten food, appears dehydrated (skin tents e.g. the skin stays up when pinced)

